# A joint analysis of longevity and age-related disease variants for gene expression association

**DOI:** 10.1101/2021.01.25.428082

**Authors:** Lu Zeng, Shouneng Peng, Seungsoo Kim, Jun Zhu, Bin Zhang, Yousin Suh, Zhidong Tu

**Affiliations:** Department of Genetics and Genomic Sciences, Icahn School of Medicine at Mount Sinai, NY; Department of Obstetrics and Gynecology, Columbia University, NY

## Abstract

A large number of genetic variants associated with human longevity have been reported but how they play their functions remains elusive. We performed an integrative analysis on 113 genome-wide significant longevity and 14,529 age-related disease variants in the context of putative gene expression regulation. We found that most of the longevity allele types were different from the genotype of disease alleles when they were localized at the same chromosomal positions. Longevity variants were about eight times more likely to be associated with gene expression than randomly selected variants. The directions of the gene expression association were more likely to be opposite between longevity and disease variants when the association occurred to the same gene. Many longevity variants likely function through down-regulating inflammatory response and up-regulating healthy lipid metabolisms. In conclusion, this work helps to elucidate the potential mechanisms of longevity variants for follow-up studies to discover methods to extend human healthspan.

## Introduction

Longevity is a heritable trait and it was estimated that genetic factors account for approximately 20% to 30% of the variation in lifespan from twin studies (Herskind et al., 1996; Mitchell et al., 2001). Although a very limited number of genetic factors have been replicated for association with longevity, e.g., the ones near *APOE* and *FOXO3*, it is widely believed that longevity is influenced by a large number of variants (Pilling et al., 2016) with most of them playing a small effect on lifespan. Currently, hundreds of longevity variants have been reported (Melzer et al., 2020) from association studies that were conducted on increasingly greater population sizes. Validating these variants, elucidating their functions, and eventually translating the key findings into actionable interventions have become an important but challenging task for human aging research.

In addition to elucidating the determinant mechanisms of human lifespan, research on longevity could also shed light on the genetic regulation of age-related diseases. The relationship between longevity and age-related diseases is complex and remains to be fully elucidated. Although in general, long-lived individuals are healthier than their age-matched peers of average lifespan (Engberg et al., 2009), they are not free of age-related diseases. Actually, half of the centenarians age with at least one chronic disease and about one-fourth with disability (Ailshire et al., 2015). It has been reported that individuals with long lifespan do not have significantly lower frequencies of disease alleles (Beekman et al., 2010). Therefore, further identifying and investigating the longevity variants that can prevent or slow the development of age-related diseases will be of great importance for healthy aging research.

One strategy to study longevity variants is to experimentally interrogate the function of each variant (Flachsbart et al., 2017); although such approach is critical, it is time consuming and does not scale-up easily. It has been suggested that integrative data-driven approach may help us to gain new insights into the validity and functionality of the longevity variants. For example, Dato et al. performed a pathway-based SNP-SNP interaction analysis to investigate beyond the effect of single SNP (Dato et al., 2018). In this work, we use data from GTEx (Genotype-Tissue Expression) project to portray the landscape of genetic association with gene expression by jointly considering longevity and age-related disease variants. In particular, we ask a few key questions, do longevity variants represent a distinct set of genetic factors from disease variants? What fraction of longevity variants are likely to function through regulating gene expression in human? And finally, when longevity and disease variant are both associated with a gene’s expression, will the associations be in the same or different directions? We believe addressing these questions will help us to better understand longevity variants and their relationship with disease variants.

## Results

### Longevity alleles are mostly different from disease alleles when they are localized at the same chromosomal positions

To obtain a comprehensive list of longevity variants, we relied on three data sources, i.e., the NHGRI-EBI GWAS Catalog (Buniello et al., 2019), LongevityMap (Budovsky et al., 2013) and manually curated published studies, encompassing 90 GWA studies published from 1991 to 2019 (Fig. 1). In total, 3,674 genetic loci associated with longevity were collected from these data sources. Among them, 113 longevity variants were genome-wide significant (p≤5×10^−8^), and 104 of them were genotyped in the GTEx v8 data which contained genotype information for 46,569,704 SNPs (Supplemental Table S1).

**Figure 1.**
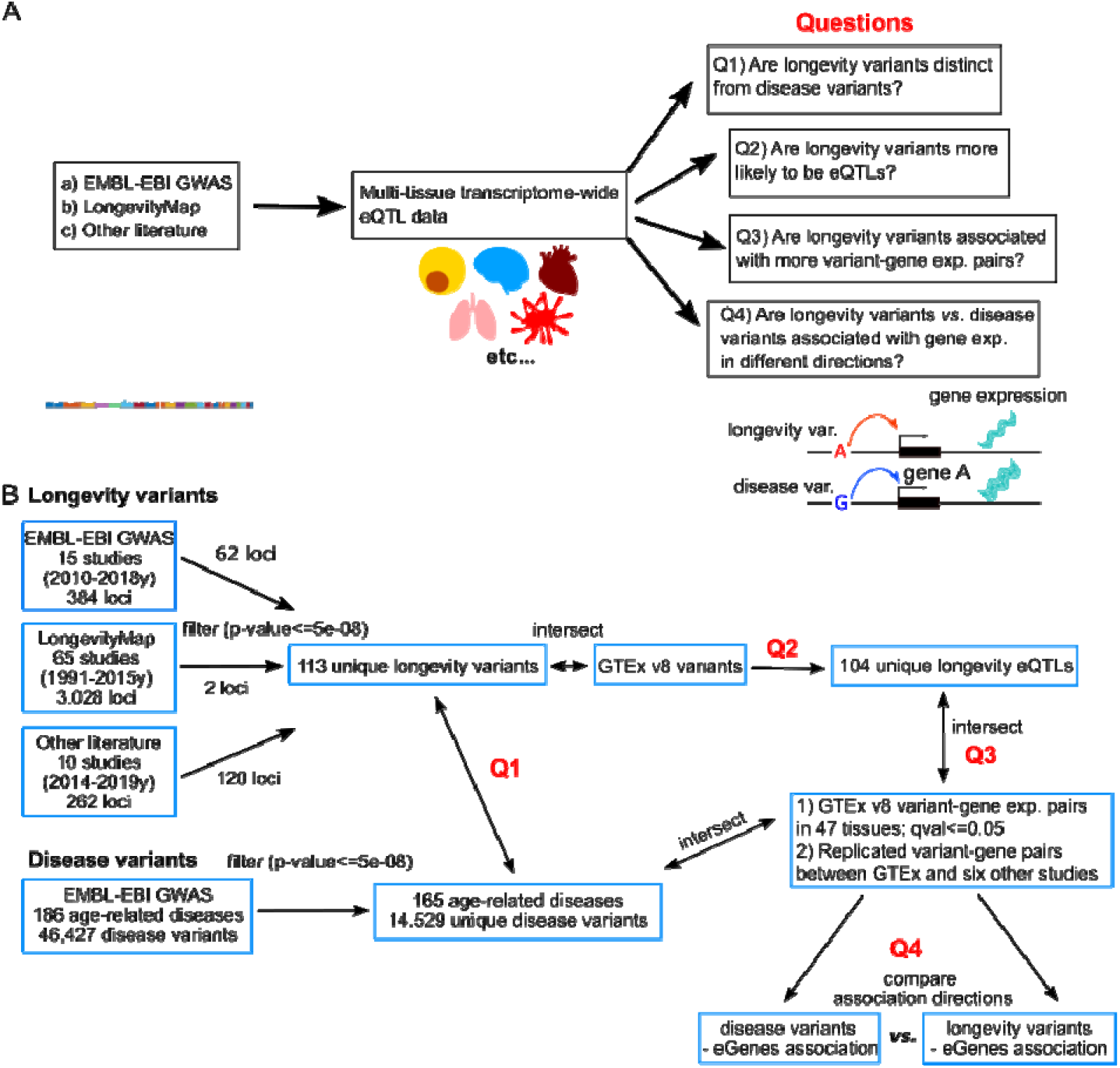
A. An overflow of the study. We highlight the key questions which we address by comparing longevity and disease variants in the context of gene expression association. B. Additional information on data input and the analytical pipeline, questions Q1 to Q4 correspond to the questions in A.

We next investigated if the loci for these 113 longevity variants could also be associated with age-related diseases. To do so, we downloaded genetic variants associated with 165 age-related disease categories whose incidence rates increase exponentially with age (Jaul & Barron, 2017) from the NHGRI-EBI GWAS Catalog, covering diseases such as Alzheimer’s disease, type 2 diabetes, hypertension and cancers. Similar to longevity variants, we only retained disease variants that were genome-wide significant (p≤5×10^−8^), we then removed disease variants whose risk alleles were inconsistent within the same disease category (e.g., rs11257238 alleles T and C were both annotated to increase the risk of Alzheimer’s disease and were removed due to such inconsistency (Jansen et al., 2019; Jun et al., 2017)), which resulted in a total number of 14,529 unique disease variants (Supplemental Table S2). Our results showed that 41% (46/113) of our longevity variants were also associated with these disease traits (Support Table 1). Not surprisingly, we found that for these colocalized longevity alleles, their allele types were mostly different from the disease alleles (42 out of 46), while only four of them shared the same alleles with disease traits, namely, rs429358, rs4420638, rs16991615 and rs3184504. Since in many cases the longevity variants were identified independently from disease GWA studies which relied on different cohorts and/or examined different traits, this indicates that a large proportion of these 113 genome-wide significant longevity variants are likely real and biologically meaningful despite they have been mostly unreproducible so far (otherwise, it will be nearly impossible to observe such a high “reverse” overlap by random chance). For the four longevity variants that shared the same alleles with disease, we found that the alternative alleles of the longevity variants were often associated with some disease traits as well. rs429358, located in the fourth exon of the *APOE* gene, causes amino acid change in *APOE* protein that leads to switch from *APOE ε*3/*ε*2 to *ε*4 (Rall et al., 1982). The longevity beneficial allele T (Pilling et al., 2017; Timmers et al., 2019) is somehow associated with an increased risk for type 2 diabetes (Mahajan Taliun, et al., 2018), while the alternative allele C is associated with an increased risk of AD and AD-related measurements (such as amyloid-beta/p-tau/t-tau) (Moreno-Grau et al., 2019). rs4420638, located in the *APOC1* gene and 14kb downstream of the *APOE ε*4 allele, has a strong association with AD. It is in a strong linkage equilibrium (LD) with rs429358 (D’=0.86) (Nyholt et al., 2009). The longevity allele A (Fortney et al., 2015; McDaid et al., 2017) was found to be associated with increased age-related macular degeneration (AMD) risk (Fritsche et al., 2013); while the alternative allele G was associated with increased risk of LDL, CAD, AD, and higher all-cause mortality (Deelen et al., 2014). rs16991615 is a nonsynonymous SNP on chromosome 20. The longevity allele A (Bae et al., 2019) was also associated with increased risk of breast cancer (Michailidou et al., 2017) and heavy menstrual bleeding (Gallagher et al., 2019). rs3184504 is a nonsynonymous SNP in the *SH2B3* gene. The longevity allele C (Pilling et al., 2017) was associated with an increased risk of colorectal cancer while the alternative allele T was associated with an increased risk of CAD (Schunkert et al., 2011) and stroke (Malik et al., 2018).

Among the 46 colocalized variants, 37 (>80%) were eQTLs in the GTEx data, while only 7.8% of randomly selected GTEx SNPs were eQTLs (see details in the next section), indicating longevity variants co-localized with disease traits are highly enriched for loci associated with gene expression.

### Longevity variants are significantly more likely to be associated with gene expression

In GTEx data, 67 longevity variants were associated with 183 unique genes’ expression (also known as eQTL-eGene pairs) in 47 GTEx tissues (false discovery rate (FDR) ≤0.05), which corresponded to a total of 1,793 eQTL-eGene pairs. This reduced to 326 unique variant-gene pairs if we did not consider tissue specificity, i.e., by counting the same eQTL-eGene pair across different tissues as one pair (Support Table 2 & Supplemental Table 3).

Permutation test was used to assess if the observed number of longevity eQTLs would occur by random chance. Since 104 of 113 longevity variants were genotyped in the GTEx v8 data, we randomly selected 104 variants from all the 46,569,704 GTEx SNPs for 1,000 times (without replacement) and counted how many of them were eQTLs. Compared to 67 out of 104 longevity variants that were eQTLs, in the permutation runs, on average, 8.4 randomly selected variants were eQTLs, with a standard deviation (SD) of 3.34. This indicates that longevity variants are significantly more likely to be associated with gene expression compared to random SNPs (Fig. 2A, permutation test, mean=8.4, SD=3.34, p≤1.0×10^−3^).

**Figure 2.**
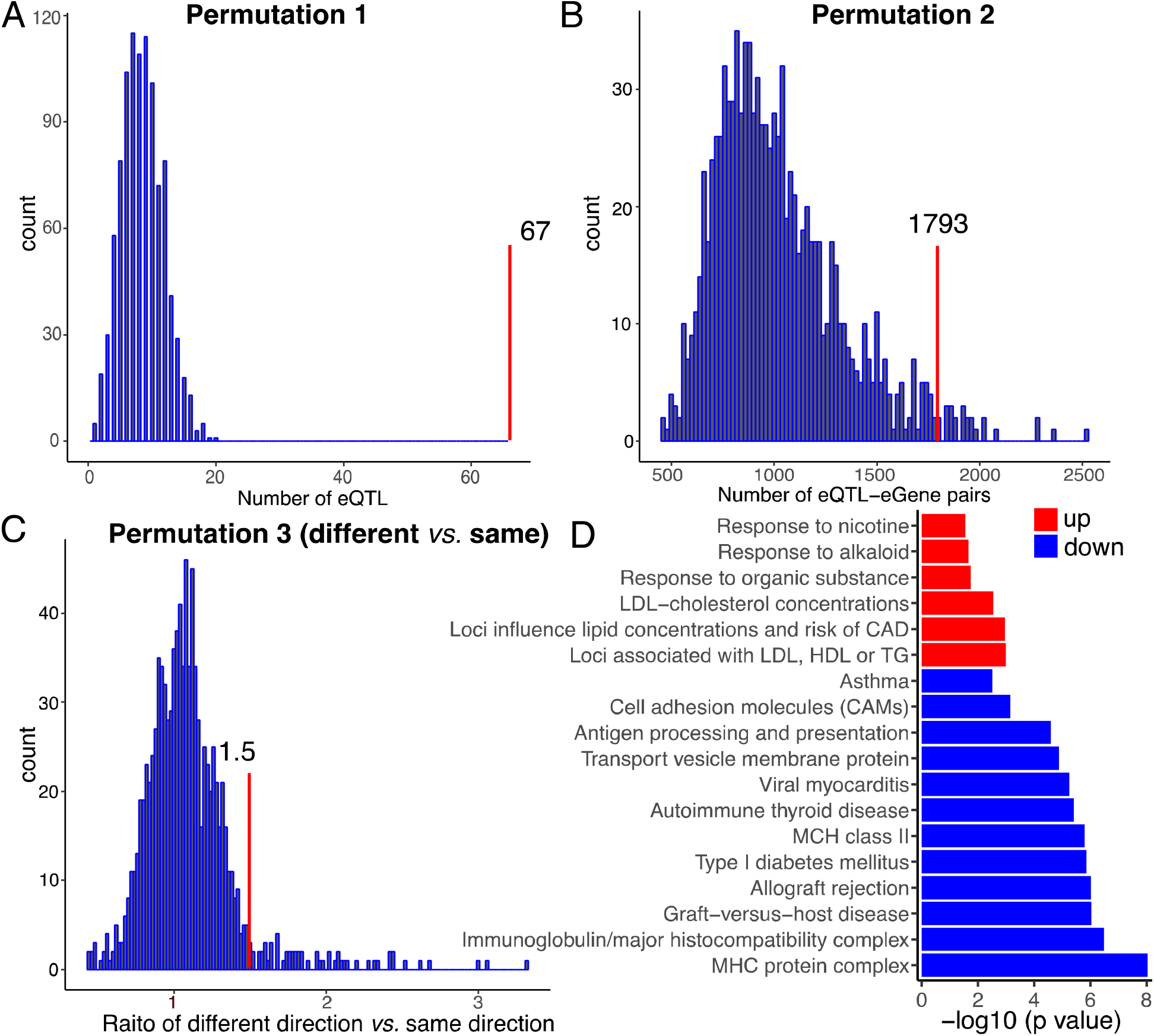
Permutation tests were performed to evaluate (a) if longevity variants were more likely to be eQTLs compared with randomly selected SNPs, (b) if longevity variants were involved in more variant-gene expression association pairs compared with randomly selected eQTLs, (c) the 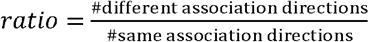 indicates if longevity variants were more likely to associate with gene expression in the opposite direction from disease alleles when they both were associated with same gene expression in subcutaneous fat, and (d) functional enrichment of longevity-variants associated eGenes.

Similarly, to estimate the significance of the observed number of longevity variant-gene expression pairs, we randomly selected 67 GTEx eQTLs for 1,000 times, and calculated the number of variant-gene expression pairs for these eQTLs across 47 GTEx tissues. The permutation analysis revealed that 67 randomly selected eQTLs were involved in an average of 1021.4 variant-gene expression pairs (SD=311.90), compared to 1,793 longevity variant-gene pairs (empirical p=0.025). When we did not consider the tissue specificity of the eQTLs, the permutation runs identified 198.1±30.68 (mean±SD) variant-gene pairs, which is significantly less than the observed 326 longevity variant-gene pairs (empirical p≤1.0×10^−3^). This result suggests that longevity eQTLs were significantly associated with more eGenes in GTEx than randomly selected eQTLs (Fig. 2B).

### Longevity variants associated up- and down-regulated eGenes were involved in distinct functions

We next investigated the function of eGenes that were associated with longevity variants. We only focused on eQTLs showing relatively consistent association directions with gene expression across GTEx tissues and we considered up- and down-regulated eGenes separately. In particular, we kept eQTL-eGene pairs if their directions were consistent across ≥ 60% GTEx tissues, and the expression direction of these eGenes were determined by the majority directions (≥ 60%). For instance, longevity variant rs72738736 allele G (Pilling et al., 2017) showed opposite association with iron responsive element binding protein 2 (*IREB2*) gene expression in two tissues: whole blood (higher gene expression) and testis (lower gene expression) (Support Table 2). Since there is no simply way to determine which associations are biologically meaningful, we removed this eQTL-eGene pair for further analysis and only focus on those eQTLs showing an apparent dominant association direction.

Based on GTEx significant variant-gene expression associations (FDR≤0.05), a slightly higher proportion of the longevity alleles were associated with increased gene expression. For example, 67 longevity eQTLs were found associated with 1,751 variant-gene pairs, and 944 of them were associated with higher gene expression with longevity alleles, corresponding to 94 unique genes, while 807 of them were down-associated with longevity alleles, corresponding to 89 unique genes. When we did not count the tissue specificity, the total number of variant-gene pairs reduced to 317 and among them, 167 were up, and 150 were down for the association with longevity alleles.

We then inspected the function enrichment for genes associated with longevity variants using DAVID tools (Huang da et al., 2009). GO annotation and pathway analysis demonstrated differential function enrichment between up- and down-associated genes with longevity variants. Among genes down-associated with longevity variants, they were characterized for MHC protein complex (Benjamini-Hochberg (BH)=9.19×10^−9^), immunoglobulin (BH=3.23×10^−7^) and antigen processing (BH=2.59×10^−5^). On the other hand, genes up-associated with longevity variants were enriched for functions such as response to blood low-density lipoprotein cholesterol, high-density lipoprotein or triglycerides in humans (BH=1.03×10^−3^), and newly identified loci that influence lipid concentrations and risk of coronary artery disease (BH=1.07×10^−3^) (Figure 2D, a full list of functional annotations is provided in Supplemental Table S4). This result suggests that longevity variants could play their beneficial roles through down-regulating the inflammatory response and up-regulating healthy lipid metabolisms.

### The association directions were more likely to be opposite when longevity and disease variants were associated with same gene’s expression

We wanted to obtain a global view of the potential gene expression regulation by longevity variants and compare that with 14,529 unique age-related disease variants as aforementioned.

To examine the potential influence of longevity and disease variants on gene expression, we focused on eGenes whose expression was simultaneously associated with at least one disease and one longevity variant in GTEx data. Using GTEx subcutaneous fat tissue as an example, 30 longevity variants were associated with 60 genes’ expression, corresponding to 92 longevity variant-gene expression pairs. 360 unique disease variants from 76 disease traits were also associated with these 60 genes’ expression, forming a total of 1,513 unique longevity variant-gene expression-disease variant trios. We then inspected the association directions between the gene expression of these 60 genes with disease *vs*. longevity variants. The comparison showed that the direction of association with gene expression for longevity alleles were different from disease alleles for 931 unique variant-gene-variant trios. For example, the longevity beneficial allele G of rs3130507 (Pilling et al., 2017) was associated with lower expression of transcription factor 19 (*TCF19*), while type 2 diabetes variants rs2073721 (Mahajan Wessel, et al., 2018) was associated with an increased *TCF19* gene expression in GTEx adipose tissue. *TCF19*’s mRNA expression has been reported to increase in nondiabetic obesity (Krautkramer et al., 2013). In contrast, 621 unique variant-gene-variant trios showed same gene expression association direction with longevity *vs*. disease alleles. For example, as we mentioned above, longevity variants rs3130507 was associated with down-regulated gene expression of *TCF19*, while the allele G of rs3094013, a SNP associated with increased body mass index (Graff et al., 2017) was also associated with decreased *TCF19* gene expression in GTEx adipose tissue.

For nine genes, their gene expression associations with longevity *vs*. disease alleles were consistently in the opposite direction in GTEx subcutaneous fat. These include FES proto-oncogene, tyrosine kinase (*FES*), neutrophil cytosolic factor 1C pseudogene (*NCF1C*), NOP2/sun RNA methyltransferase 5 pseudogene 1 (*NSUN5P1*), PMS1 homolog 2, mismatch repair system component pseudogene 2 (*PMS2P2*), PMS1 homolog 2, mismatch repair system component pseudogene 3 (*PMS2P3*), POM121 transmembrane nucleoporin C (*POM121C*), speedy/RINGO cell cycle regulator family member E5 (*SPDYE5*), and stromal antigen 3-like 1 (pseudogene) (*STAG3L1*). Seven of these genes showed decreased gene expression association with longevity alleles, except for *FES* and *PMS2P3* which showed increased gene expression. The *FES* expression was down-regulated in colorectal cancer cell lines (Shaffer & Smithgall, 2009).

Similarly, we performed a permutation test to evaluate the significance of observing more opposite associations with gene expression between longevity and disease variants. Since 30 longevity variants were involved in 92 variant-gene pairs in subcutaneous fat, we randomly selected 30 eQTLs for 1,000 times from GTEx subcutaneous adipose tissue while we ensured that these variants were involved in 92 variant-gene pairs in each run (see details in Methods). We then extracted GTEx adipose eGenes associated with these 30 eQTLs to pull out all the variants that were associated with these eGenes; by overlapping these variants with disease variants from the 165 disease categories, we then checked the direction of association between disease alleles and longevity alleles for the corresponding eGenes. From this permutation analysis, we observed a slightly more variant-gene expression-variant trios in the opposite association directions, as the mean value of the ratio between opposite direction of association (506.2±189.82) *vs*. same direction of association (471.9±177.25) was 1.1 with a SD of 0.23. In the real data, this ratio was 1.5 (931 associations in opposite directions between longevity *vs*. disease alleles, and 621 associations in the same direction), suggesting longevity and disease variants are significantly more likely to associate with a gene’s expression in the opposite directions (empirical p=0.042).

Next, we expanded this analysis to all the other 46 GTEx tissues; we found that 3,148 unique longevity variant-gene expression-disease variant trios showed opposite associations between longevity and disease alleles, while 1,971 unique longevity variant-gene expression-disease variant trios had same association direction, leading to a ratio of 1.6. Twenty eGenes consistently showed opposite gene expression association between longevity and disease alleles (Table 1), including two genes (*NSUN5P1* and *POM121C*) identified from subcutaneous fat and 18 other genes, which are *AP4B1-AS1, AREL1, CIB4, FURIN, CASTOR2, GULOP, KCNK3, KLHDC10, PBX2, PHTF1, PMS2, PMS2CL, RGS12, SLC22A1, SPDYE12P, TOMM40, USP28* and *ZC3HC1* (detailed information can be found in support table 3).

**Table 1.** List of genes whose expression associated with longevity and disease variants in consistent opposite directions across 47 GTEx tissues. We identified twenty genes whose association of their gene expression with longevity and disease alleles were consistently in the opposite directions. The third column shows the longevity eQTL for each gene and the beneficial allele. The fourth column shows the association direction with the longevity variant. The fifth column denotes the disease eQTLs and the risk alleles; the sixth column is the association direction with the disease variant; the last column shows the number of GTEx tissues in which such opposite associations were observed.

| Gene symbol | Gene name | Longevity variant and allele type | Exp dir | Disease variant and allele type | Exp dir | No. of GTEx tissues |
| --- | --- | --- | --- | --- | --- | --- |
| *AP4B1-AS1* | Adaptor related protein complex 4 subunit beta 1 antisense RNA 1 | rs1230666:G | higher | rs1230666:A<br>rs7513707: A<br>rs11552449:T | lower | 2 |
| *AREL1* | Apoptosis resistant E3 ubiquitin protein ligase 1 | rs61978928:C | lower | rs2165197:C | higher | 1 |
| *CIB4* | Calcium and integrin binding family member 4 | rs1275922:G | lower | rs1275984:A<br>rs1275982:C<br>rs1731249:T<br>rs1275978:C<br>rs1731243:C<br>rs1275988:C<br>rs11126666:A | higher | 1 |
| *FURIN* | Furin, paired basic amino acid cleaving enzyme | rs6224:G | lower | rs17514846:A<br>rs2071382:T<br>rs2521501:T<br>rs4932371:C<br>rs4932373:C<br>rs8039305:C | higher | 3 |
| *CASTOR2* | Cytosolic arginine sensor for MTORC1 subunit 2 | rs113160991:G | higher | rs6944634:G | lower | 1 |
| *GULOP* | Gulonolactone L- oxidase, pseudogene | rs7844965:G | higher | rs2279590:C<br>rs11136000:C<br>rs4236673:G<br>rs1532278:C | lower | 1 |
| *KCNK3* | Potassium two pore domain channel subfamily K member 3 | rs1275922:G | higher | rs11126666:A<br>rs1275978:C<br>rs1275982:A<br>rs1275984:A<br>rs1275988:C<br>rs1731243:C<br>rs1731249:T | lower | 3 |
| *KLHDC10* | Kelch domain containing 10 | rs56179563:A | higher | rs11556924:C | lower | 1 |
| *NSUN5P1* | NOP2/Sun RNA methyltransferase 5 pseudogene 1 | rs113160991:G | lower | rs6963105:G<br>rs6944634:G<br>rs1167821:T<br>rs1167827:G<br>rs1167796:G | higher | 36 |
| *PBX2* | PBX homeobox 2 | rs28383322:T | lower | rs2071288:C<br>rs9268856:C | higher | 1 |
|  |  |  |  | rs41268896:A |  |  |
| *PHTF1* | Putative homeodomain transcription factor 1 | rs1230666:G | higher | rs1230666:A<br>rs7513707:A<br>rs11552449:T | lower | 3 |
| *PMS2* | PMS1 homolog 2, mismatch repair system component | rs1830074:C | lower | rs1830074:C<br>rs7456039:C | higher | 1 |
| *PMS2CL* | PMS2 C-terminal like pseudogene | rs3764814 :C | higher | rs1830074 :C | lower | 2 |
| *POM121C* | POM121 transmembrane nucleoporin C | rs113160991:G | lower | rs6963105:G<br>rs6944634:G<br>rs1167821:T<br>rs1167827:G<br>rs1167796:G<br>rs34324971:A | higher | 24 |
| *RGS12* | Regulator of G protein signaling 12 | rs61348208:T | higher | rs362275:C<br>rs3121419:C<br>rs363066:T | lower | 2 |
| *SLC22A1* | Solute carrier family 22 member 1 | rs1510224:T<br>rs111333005:G | higher | rs3798220:C<br>rs9295128:T<br>rs140570886:C<br>rs186696265:T<br>rs2297374:C | lower | 7 |
| *SPDYE12P* | Speedy/RINGO cell cycle regulator family member E12, pseudogene | rs113160991:G | lower | rs35005436:C<br>rs6944634:G | higher | 1 |
| *TOMM40* | Translocase of outer mitochondrial membrane 40 | rs71352238:T | lower | rs34342646:A<br>rs71352238:C<br>rs10119:A | higher | 1 |
| *USP28* | Ubiquitin specific peptidase 28 | rs61905747:A | higher | rs61904987:T | lower | 1 |
| *ZC3HC1* | Zinc finger C3HC-type containing 1 | rs56179563:A | lower | rs11556924:C | higher | 2 |

### Longevity eQTLs from GTEx were replicable in independent studies

To evaluate the robustness of GTEx eQTLs, we collected eQTL-eGene pairs from six independent eQTL studies which covered five tissues: adipose (MUTHER) (Nica et al., 2011), brain cortex (ROSMAP) (Ng et al., 2017), heart left ventricle (Koopmann et al., 2014), lung (Hao et al., 2012) and two blood studies (Võsa et al., 2018; Westra et al., 2013). We compared eQTLs from these independent studies with the eQTLs identified in the corresponding GTEx tissues. The comparison showed that eQTLs from these studies were highly enriched in GTEx data. For example, 37% adipose eQTLs from the MUTHER study were reproducible in GTEx subcutaneous fat; for two blood eQTL studies, 44% of the eQTLs from Võsa’s work and 61% of the blood eQTLs from Westra’s work matched with GTEx whole blood eQTLs. For eQTL-eGene pairs, a tissue-specific enrichment pattern could be seen for each tissue. For example, GTEx subcutaneous fat had the most matching eQTL-eGene pairs with the adipose MUTHER study (>2,000 pairs), while it had much fewer pairs in GTEx brain cortex (893 pairs). Similarly, 4,278 eQTL-eGene pairs from the independent lung eQTL study were found replicated in the GTEx lung, but much fewer pairs were found in other tissues (Supplemental Table S5). In addition, we found most of the reproducible variant-gene pairs showed same direction in eQTL-gene expression between independent studies and GTEx (Supplemental Table S5). For instance, in blood tissue, 1,073,253 variant-gene pairs showed the same direction in eQTL-gene expression association between GTEx and independent studies, only 123,723 pairs showed different directions. For the following analysis, we only considered the variant-gene expression pairs that showed the same association direction between GTEx and those independent studies.

We then repeated part of our analyses with these replicated eQTL-eGene pairs. Our results showed that overall the eQTL-gene expression association directions were different between disease and longevity alleles in all five tissues, resulting 531 unique variant-gene-variant trios showed opposite directions, and 203 unique variant-gene-variant trios showed same directions (Supplemental Tables S6 & 7), a ratio of 2.6 between the two. Since this ratio increased from 1.5 in subcutaneous fat, to 1.6 across 47 tissues, to 2.6 in replicated eQTL studies, this indicates that the longevity and disease alleles are more likely to associate with gene expression in the opposite directions when robust eQTLs are considered.

For these replicated eQTL-eGene pairs, we noticed that longevity alleles showed consistent opposite association compared to disease alleles on eleven genes’ expression, five of them were replicated from our previous analysis (*NCF1C, NSUN5P1, PMS2P3, RGS12* and *SLC22A1*), and the rest are *CHRNA5, HLA-B, MAPKAPK5, POU5F1, PBX2* and *RNF5*. In several cases, longevity alleles were found to play a putative beneficial role with respect to the associated gene expression (Table 2). For examples, longevity variant rs61348208 (Timmers et al., 2019) was associated with higher *RGS12* gene expression, while its expression level was relatively lower in African American (AA) prostate cancer (Wang et al., 2017). Longevity variants rs3130507, rs1510224 and rs186696265 (Pilling et al., 2017) were found to be associated with higher *SLC22A1* gene expression, while the expression of *SLC22A1* was down-regulated in hepatocellular carcinoma (Okabe et al., 2001). Lower *HLA-B* gene expression was associated with longevity variants rs3131621 and rs3130507 (Pilling et al., 2017), while the down-regulation of *HLA-B*/*C* expression has been reported to correlate with a lower tumor stage and a longer disease-free survival in colorectal cancer patients (Menon et al., 2002).

**Table 2.**
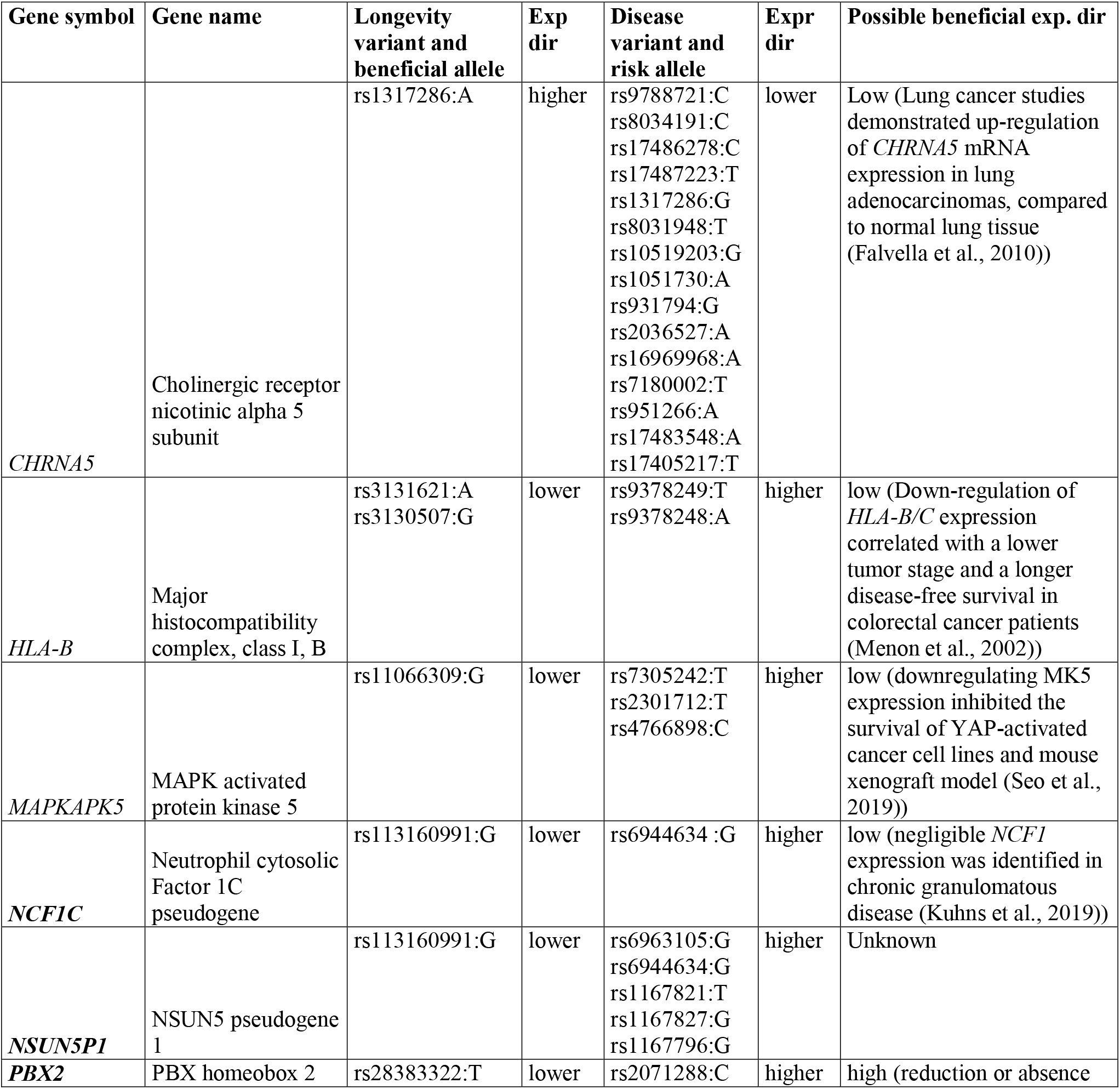
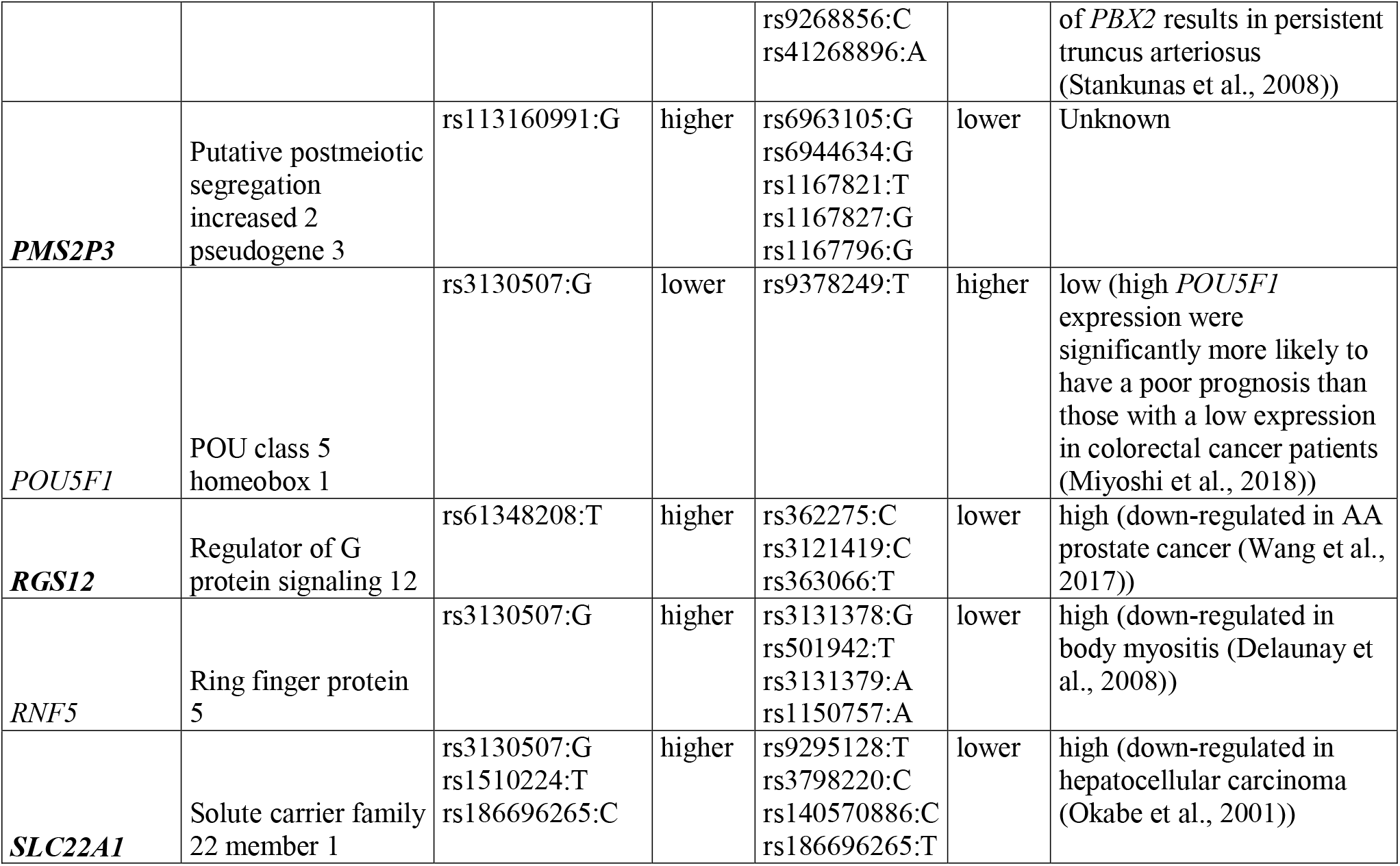
Genes whose expression was associated with longevity vs. disease variants in opposite directions in multiple eQTL studies. We identified eleven genes whose association of their gene expression with longevity and disease alleles were consistently in the opposite directions (gene symbols in bold-fonts are those replicated in GTEx data). The fourth column “Exp dir” shows the association direction of longevity alleles with each gene’s expression; the last column is the putative beneficial expression direction of the corresponding genes based on previous studies, “unknown” indicates that no related expression studies were found for the corresponding gene.

## Discussion

Thousands of genetic variants have been reported to be associated with human longevity (Budovsky et al., 2013). Validation and follow-up studies on such a large number of longevity variants remain to be challenging. For example, many of the variants are located in the non-coding or intergenic regions and do not pin-point to a protein coding gene, how they may play beneficial functions (if any) to human lifespan is elusive.

In this study, we collected 113 genome-wide significant longevity variants, and investigated their relationship with age-related disease variants in the context of gene expression regulation. It is of note that heterogeneous study designs were taken to identify the genetic determinants of human longevity, e.g., population-based case and control cohorts (Fortney et al., 2015; Zeng et al., 2016), family-based cohorts to study parental survival trait (Joshi et al., 2017; Pilling et al., 2017) and prospective studies (Broer et al., 2015; McDaid et al., 2017). Therefore, the longevity variants compiled from these studies may point to different biology of aging and lifespan, even though we collectively call them longevity variants. The heterogeneity in the derivation of longevity variants could potentially complicate the interpretation of our results.

We made several interesting observations in this work: first, from longevity GWA studies, not every hit may be beneficial for longevity, particularly if we are looking for variants responsible for healthspan. This is because some age-related diseases could be enriched in very long-lived individuals, such as the AMD. Therefore, it is possible that some of the GWAS derived longevity variants are causally associated with these diseases and represent increased disease risks although they are also associated with longer lifespan. Second, a large proportion of the genome-wide significant longevity variants may represent true biological signals even though they have been hardly replicated. This is supported by the fact that over 80% of the longevity variants when colocalized with disease variants had different allele types from the disease risk alleles. Since disease and longevity variants were often identified from independent studies and derived based on very different traits, this suggests that many of the genome-wide significant longevity variants are not random hits but convey biological signals. This is expected as long-lived people are less likely to develop age-related chronic conditions than their same-age peers, and they remain healthier longer into the old ages (Sebastiani et al., 2017; Wei et al., 2017). Third, we think that many longevity variants could play their functions through modulating underlying gene expression. This is supported by the observation that longevity variants were about eight times more likely to be eQTLs in the GTEx data compared with randomly selected SNPs. In particular, we observed that when a gene expression is associated with both longevity and disease variants, the association directions are more likely to be different than being the same. This trend is more apparent when we narrowed down to those reproducible eQTL-eGene pairs from independent studies. As a key goal of human aging and geroscience research is to identify methods to promote human healthspan, this joint analysis provides a data-driven approach to prioritize the longevity variants that may function through counteracting the effect of disease variants through gene regulation.

Although we observed that longevity and disease variants were often associated with a gene expression in opposite directions, many of them also showed same direction in association with gene expression. This could receive multiple explanations. First, reported genetic variants for longevity and diseases may not always be accurate; in fact, some of the reported findings on these variants were inconsistent. For example, allele T of rs660240 was associated with increased heel bone mineral density based on the NHGRI-EBI GWAS Catalog, while it was also reported to associate with increased risk of osteoporosis in certain studies (Morris et al., 2019), while low bone mineral density (BMD) is the major determinant for osteopenia and osteoporosis (Cauley et al., 2007). Currently we largely relied on the GWAS catalog annotation since manually curating 14,529 SNPs will require an extraordinary amount of effort and in some cases impossible without full access to the original genotype/phenotype data; Second, some eQTL calls may not be robust; although we show they are largely consistent across different studies, there were some apparent variations in eQTL calling across studies; Third, eQTLs only indicate association between genetic variants and gene expression, it remains unknown if the eQTLs are actually involved in regulating the gene expression; this becomes even more complicated due to the fact that multiple mapping among variants-gene expression exists. For instance, rs602633 (Pilling et al., 2017) was associated with the *PSRC1, SYPL2, CELSR2, SORT1* expression changes across 25 GTEx tissues, it is unclear which variant-gene expression association is biologically functional; last but not least, since it is not known for certain in which tissue(s) a variant plays its function, we may consider eQTLs that are actually not functional in the tissues under consideration. Recently, multiple statistical methods have been developed to facilitate the casual inference between genetic variant, gene expression and phenotypic outcome. For example, the COLOC is a Bayesian test to assess if two traits shared a causal variant (Giambartolomei et al., 2014), S-PrediXcan was developed to perform transcriptome-wide genome association to identify possible gene expression associated with the phenotypic traits (Jasinska, 2020), and transcriptome-wide summary statistics-based Mendelian randomization method TWMR was developed to use multiple SNPs as instruments and multiple gene expression traits as exposures to infer if the causal link between instruments and outcome was mediated by exposures (Porcu et al., 2019). Although these methods are helpful to test the potential involvement of gene expression in causing phenotypic trait, they all require GWAS summary statistics and therefore limited their use in this study.

We noticed that two well-known longevity genes *APOE* and *FOXO3* were not included in our gene list as shown in Tables 1 and 2. Two common *APOE* alleles were found either significantly depleted (*ε*4 allele) or enriched (*ε*2 allele) in long-lived individuals as compared to controls (Ryu et al., 2016). The variants for both alleles are located in protein-coding regions and may not affect regulatory regions to impact gene expression. In fact, we found three of our longevity variants (rs283811 (Sebastiani et al., 2017), rs7412 (Deelen et al., 2019) and rs4420638 (McDaid et al., 2017)) were associated with *APOE*’s gene expression only in GTEx skin tissues but not in other tissues. Although *FOXO3* has been identified to associate with longevity in human, none of the reported SNPs reached genome-wide significant (p≤5×10^−8^) in our initial longevity variants collection, therefore the longevity variants associated with *FOXO3* were filtered out (Flachsbart et al., 2009; Li et al., 2009; Willcox et al., 2008). In addition, none of our longevity variants were significantly associated with *FOXO3*’s gene expression in GTEx data.

Longevity is believed to be a highly polygenic phenotype and we have only examined a small fraction of all possible longevity variants as we focused on genome-wide significant variants in this study. We expect to see more longevity variants to emerge and to be replicated as genetic information will be increasingly available from larger populations. This work therefore establishes a new direction for further exploration to prioritize the longevity variants that could be truly beneficial to extending human health span.

## Methods

### Longevity and disease variants collection

In order to get a comprehensive list of longevity variants, we went through three sources: 1) LongevityMap: a database of human genetic variants associated with longevity, including 65 GWA studies carried out from 1991 to 2015, with 3,028 loci collected; 2) The NHGRI-EBI GWAS Catalog: a curated collection of human genome-wide association studies, including 15 longevity GWAS studies with 384 loci; 3) last but not least, by going through recent longevity studies that were not covered by either LongevityMap or the NHGRI-EBI GWAS Catalog. We manually curated 262 longevity variants (Fig. 1). We only considered longevity variants with a genome-wide significant p-value (≤5×10^−8^), which resulted to 113 longevity variants with reported beneficial alleles (Supplemental Table S1). 104 of these 113 longevity variants were covered in GTEx genotype data. The 104 longevity variants were then intersected with 68,129,832 significant variant-gene expression associations (FDR≤0.05) identified in GTEx v8. 67 longevity variants were found associated with 183 unique genes’ expression, corresponding to 1,793 variant-gene pairs across 47 tissues in a tissue-specific manner.

Genetic variant information for 186 age-related diseases were downloaded from the NHGRI-EBI Catalog (accessed on July 10, 2019), only variants reaching genome-wide significance (p≤5×10^−8^) with known risk alleles were kept. We removed disease variants whose risk alleles were inconsistent within the same disease category. After filtering, a total of 14,529 unique variants with respect to 165 disease traits were compiled for further analysis.

### Permutation test

Permutation analyses were carried out to: 1) test if longevity variant is more likely to be an eQTL; we randomly selected 104 variants from all SNPs genotyped in GTEx for 1,000 times, and we checked for each run how many SNPs were associated with gene expression. Empirical p-value was then calculated to estimate the significance of the observed number of eQTLs for longevity variants; 2) test if longevity variants were involved in more variant-gene pairs than randomly selected variants; we randomly selected 67 variants that were eQTLs in GTEx for 1,000 times, and counted how many variant-gene expression pairs were linked to these eQTLs in GTEx, similarly, empirical p-value was then calculated to test the significance of observed eQTL-eGene pairs for longevity variants; 3) finally, we tested the significance of the difference between longevity *vs*. disease alleles on gene expression association direction. We used subcutaneous fat as an example, 30 unique longevity variants were found to be associated with 60 genes’ expression, corresponding to 92 variant-gene pairs. To perform the permutation test, we first randomly selected 30 GTEx eQTLs for 1,000 times that had same structures as longevity eQTLs in subcutaneous fat. For example, 15 longevity variants were found to be associated with only one gene’s expression, while 6 longevity variants were associated with two genes’ expression in the adipose tissue and so on. GTEx eQTLs were thus randomly selected accordingly based on the same structure in subcutaneous fat, i.e., to randomly select 15 eQTLs that were associated with only one gene’s expression, and 6 eQTLs that were associated with two genes’ expression, etc. In addition, 20 out of 30 longevity beneficial alleles were major alleles for GTEx adipose, and 10 were minor alleles. We then randomly selected 20 major alleles and 10 minor alleles as longevity beneficial alleles in each random eQTL set.

### Cross-studies eQTL comparison

Six independent eQTL studies were downloaded to evaluate the robustness of GTEx eQTLs, covering five tissues: adipose (Nica et al., 2011), brain cortex (Ng et al., 2017), left ventricle (Koopmann et al., 2014), lung (Hao et al., 2012) and two blood studies (Võsa et al., 2018; Westra et al., 2013). The reproducible variant-gene pairs between these studies and the corresponding GTEx tissues were then counted based on three criteria: 1) eQTLs from other studies were replicated in GTEx v8; 2) these replicated eQTLs associated with same eGene as in GTEx; 3) variant-gene pairs showed same association direction in eQTL-gene expression between independent studies and GTEx.

## Acknowledgements

This work was funded by NIH grant R01AG055501 to Z.T. and Y.S. The content is solely the responsibility of the authors and does not necessarily represent the official views of the National Institutes of Health. This work was also supported in part through the computational resources and staff expertise provided by Scientific Computing at the Icahn School of Medicine at Mount Sinai.

