## Supplemental files for "A joint analysis of longevity and age-related disease variants for gene expression association"

**Supplementary tables**

**Table S1. Genome-wide significance (p≤5×10-8) longevity variants from previous longevity studies.** Longevity rsID is the variant ID; Beneficial allele represents the allele associated with reduced hazards or long lifespan. Reported p-value means the p-value reported from previous studies; Candidate genes denotes the implicated genes associated with corresponding variants; PMID is the PubMed identifier number of the study source.

| Longevity rsID | Beneficial allele | Reported p-value | Candidate genes | PMID |
| --- | --- | --- | --- | --- |
| rs10198124 | G | 2.00E-08 | AC012593.1/AC013442.1 | 29030599 |
| rs10211471 | C | 2.30E-08 | GBX2/ASB18 | 30642433 |
| rs1036345 | A | 8.71E-09 | TMTC2 | 28329165 |
| rs10414043 | G | 9.12E-23 | APOC1 | 28329165 |
| rs10455872 | A | 8.50E-25 | LPA | 28748955/30642433 |
| rs10778936 | C | 2.49E-08 | TMTC2 | 28329165 |
| rs10778937 | T | 4.24E-08 | TMTC2 | 28329165 |
| rs10778938 | T | 1.98E-08 | TMTC2 | 28329165 |
| rs10778939 | A | 2.26E-08 | TMTC2 | 28329165 |
| rs10862576 | C | 1.43E-08 | TMTC2 | 28329165 |
| rs11065979 | C | 1.00E-12 | ATXN2/BRAP | 30642433 |
| rs11066309 | G | 1.00E-10 | PTPN11 | 29227965 |
| rs11115558 | T | 1.57E-08 | TMTC2 | 28329165 |
| rs11115563 | C | 1.98E-08 | TMTC2 | 28329165 |
| rs11115564 | C | 3.78E-08 | TMTC2 | 28329165 |
| rs111333005 | G | 6.60E-09 | IGF2R | 30642433 |
| rs113160991 | G | 7.50E-09 | POM121C | 30642433 |
| rs11513729 | C | 4.00E-11 | ALDH2/MAPKAPK5-AS1 | 29227965 |
| rs11531136 | C | 2.39E-08 | TMTC2 | 28329165 |
| rs11556505 | C | 4.12E-24 | TOMM40 | 28329165 |
| rs11685387 | A | 3.20E-11 | XRCC5 | 24518833 |
| rs1197774 | C | 3.60E-08 | APTX | 24518833 |
| rs1230666 | G | 6.40E-09 | MAGI3 | 30642433 |
| rs1275922 | G | 6.00E-09 | KCNK3 | 30642433 |
| rs12949468 | A | 5.00E-20 | TLK2 | 25918517 |
| rs12972156 | C | 4.13E-21 | NECTIN2 | 29227965 |
| rs12972970 | G | 3.02E-21 | NECTIN2 | 29227965 |
| rs1317286 | A | 2.00E-31 | CHRNA3 | 29227965 |
| rs13251813 | A | 2.10E-08 | WRN | 24518833 |
| rs13262617 | A | 3.00E-08 | TOX | 29227965 |
| rs1333042 | A | 2.00E-08 | CDKN2B-AS1 | 29227965 |
| rs1333045 | C | 1.77E-08 | CDKN2BAS | 28748955 |
| rs139137459 | G | 3.00E-08 | USP2-AS1 | 29227965 |
| rs1416280 | C | 5.00E-08 | GRIK2/AP002530.1/AL357139.1 | 25199915 |
| rs142158911 | A | 8.10E-09 | LDLR | 30642433 |
| rs143843429 | A | 4.00E-09 | LOC102724087 | 29227965 |
| rs146254978 | T | 5.00E-08 | FPGT/TNNI3K | 29227965 |
| rs1510224 | T | 4.00E-11 | LPAL2 | 29227965 |
| rs1556516 | G | 1.00E-20 | CDKN2B-AS1 | 29227965/30642433 |
| rs157581 | T | 1.41E-15 | TOMM40 | 28329165 |
| rs157582 | C | 2.86E-16 | TOMM40 | 28329165 |
| rs1627804 | C | 4.00E-08 | BEND3 | 29227965 |
| rs16991615 | A | 6.99E-12 | VIT | 31188284 |
| rs17514846 | C | 7.00E-10 | FURIN | 29227965 |
| rs17630235 | G | 8.00E-11 | TRAFD1/HECTD4 | 29227965 |
| rs17696736 | A | 1.00E-10 | NAA25 | 29227965 |
| rs184017 | T | 2.67E-15 | TOMM40 | 28329165 |
| rs186696265 | C | 8.00E-18 | AL109933.2-AL109933.1 | 29227965 |
| rs2020085 | T | 2.73E-08 | TMTC2 | 28329165 |
| rs2069837 | A | 2.00E-09 | IL6 | 26912274 |
| rs2075650 | A | 4.30E-24 | APOE/TOMM40 | 28329165/26677855 |
| rs2149954 | T | 2.00E-08 | EBF1/LINC02227 | 24688116 |
| rs2273500 | T | 2.00E-08 | CHRNA4 | 29227965 |
| rs2440012 | C | 4.00E-08 | ANKRD20A9P | 26912274 |
| rs283811 | A | 9.68E-14 | NECTIN2 | 28329165 |
| rs283815 | A | 4.20E-15 | NECTIN2 | 29227965 |
| rs28383322 | C | 5.00E-11 | HLA-DRB1/HLA-DQA1 | 29227965 |
| rs28926173 | G | 2.00E-08 | MC2R | 29227965 |
| rs3130507 | G | 2.00E-10 | PSORS1C3-POU5F1 | 29227965 |
| rs3131621 | A | 4.00E-08 | MICA/MICB/HCP5 | 29227965 |
| rs3184504 | T | 4.00E-08 | SH2B3/ATXN2 | 29227965 |
| rs34095326 | G | 9.78E-19 | TOMM40 | 28329165 |
| rs34342646 | G | 1.64E-21 | NECTIN2 | 29227965 |
| rs34404554 | C | 1.42E-24 | TOMM40 | 28329165 |
| rs34831921 | A | 4.00E-08 | HLA-DQA1/HLA-DRB1 | 29030599 |
| rs34967069 | T | 4.30E-09 | HLA-DQA1 | 30642433 |
| rs3764814 | C | 5.49E-15 | USP42 | 28329165 |
| rs429358 | C | 1.30E-56 | APOE/APOC1 | 29227965/29030599/30642433 |
| rs4420638 | A | 1.40E-17 | APOC1 | 28748955/26677855 |
| rs4426183 | T | 2.00E-11 | MYL2/LINC01405 | 29227965 |
| rs453843 | C | 2.71E-08 | TMTC2 | 28329165 |
| rs4639950 | C | 2.00E-16 | C1QTNF5/MFRP | 25918517 |
| rs55730499 | C | 2.00E-18 | LPA | 29227965/29030599 |
| rs56179563 | A | 5.60E-09 | ZC3HC1 | 30642433 |
| rs59007384 | G | 4.80E-15 | TOMM40 | 28329165 |
| rs602633 | T | 4.00E-09 | CELSR2/PSRC1 | 29227965 |
| rs6108784 | T | 1.00E-10 | C20orf187 | 29227965 |
| rs61348208 | T | 5.80E-09 | HTT | 30642433 |
| rs61905747 | A | 6.00E-09 | ZW10 | 29227965 |
| rs61949650 | T | 3.00E-08 | LINC00355/LGMNP1 | 29227965 |
| rs61978928 | T | 2.00E-08 | PROX2/YLPM1 | 29227965 |
| rs6224 | G | 1.30E-10 | FURIN/FES | 29227965 |
| rs6744653 | A | 7.00E-10 | TMEM18 | 30642433 |
| rs6857 | C | 1.68E-27 | APOE | 28329165 |
| rs71352238 | T | 3.58E-24 | TOMM40 | 28329165 |
| rs7137828 | C | 3.00E-14 | SH2B3/ATXN2 | 29227965 |
| rs72738736 | G | 1.00E-23 | IREB2 | 29227965 |
| rs7308896 | G | 3.03E-08 | TMTC2 | 28329165 |
| rs7309892 | C | 1.36E-08 | TMTC2 | 28329165 |
| rs7314554 | A | 4.85E-08 | TMTC2 | 28329165 |
| rs74011415 | G | 1.00E-08 | SEMA6D | 29227965 |
| rs7412 | T | 2.00E-12 | APOE | 31413261 |
| rs74444983 | T | 9.00E-09 | EXOC3L2/MARK4 | 29227965 |
| rs7676745 | A | 4.30E-08 | GPR78 | 31413261 |
| rs769449 | G | 1.08E-23 | APOE | 28329165 |
| rs7844965 | G | 8.00E-09 | GULOP | 29227965 |
| rs7961389 | C | 3.14E-08 | TMTC2 | 28329165 |
| rs7975671 | A | 4.37E-08 | TMTC2 | 28329165 |
| rs7976168 | A | 3.82E-09 | TMTC2 | 28329165 |
| rs8042849 | T | 1.60E-26 | CHRNA3/CHRNA5/HYKK | 29030599/30642433 |
| rs9295128 | G | 2.00E-11 | SLC22A2/SLC22A3 | 29227965 |
| rs951266 | G | 6.00E-14 | CHRNA5 | 28748955 |
| rs9923854 | C | 4.24E-10 | CETP | 22234866 |
| chr13:31871514 | T | 5.00E-08 | B3GALTL | 29227965 |
| rs12461964 | A | 8.00E-09 | EGLN2/CYP2A6 | 29227965 |
| rs1265564 | A | 6.00E-09 | CUX2 | 29227965 |
| rs12924886 | A | 1.40E-08 | HP | 30642433 |
| rs2229188 | C | 2.00E-08 | CYP51A1 | 25918517 |
| rs2519093 | C | 1.90E-08 | ABO | 30642433 |
| rs283812 | T | 3.60E-13 | NECTIN2 | 28329165 |
| rs4477283 | T | 1.17E-08 | CAPN9/C1orf198 | 28329217 |
| rs4970836 | G | 1.60E-09 | CELSR2/PSRC1 | 30642433 |
| rs9456508 | T | 4.00E-08 | SLC22A2 | 29227965 |
| rs10198124 | G | 2.00E-08 | AC012593.1/AC013442.1 | 29030599 |
| rs10211471 | C | 2.30E-08 | GBX2/ASB18 | 30642433 |

**Table S2. Number of genome-wide significant (p≤5×10-8) variants in 165 age-related disease traits/catogories.**

| Disease traits | No. of genome-wide significant variants |
| --- | --- |
| Abdominal aortic aneurysm | 15 |
| Acute coronary syndrome | 12 |
| Acute lymphoblastic leukemia | 30 |
| Acute myeloid leukemia | 11 |
| Adiponectin measurement | 35 |
| Age-related hearing impairment | 6 |
| Age-related macular degeneration | 553 |
| Alzheimer's disease | 266 |
| Amyotrophic lateral sclerosis | 10 |
| Aortic aneurysm | 15 |
| Apnea | 4 |
| Atherosclerosis | 63 |
| Atrial fibrillation | 306 |
| Atrophic macular degeneration | 2 |
| Back pain | 3 |
| Barrett's esophagus | 16 |
| Basal cell carcinoma | 95 |
| B-cell acute lymphoblastic leukemia | 13 |
| Benign prostatic hyperplasia | 20 |
| Beta-amyloid 1-42 measurement | 11 |
| Bladder cancer | 16 |
| Blood pressure | 1,992 |
| Body mass index | 1,809 |
| Bone density | 17 |
| Brain aneurysm | 38 |
| Breast cancer | 468 |
| Brugada syndrome | 3 |
| Cardiac arrest | 3 |
| Cardiomyopathy | 2 |
| Cardiovascular disease | 267 |
| Cataract | 6 |
| Cerebral amyloid deposition | 28 |
| Cervical cancer | 7 |
| Chronic kidney disease | 287 |
| Chronic lymphocytic leukemia | 74 |
| Chronic obstructive pulmonary disease | 448 |
| Chronic pancreatitis | 2 |
| Chronic venous insufficiency | 2 |
| Cognitive decline measurement | 42 |
| Cognitive impairment | 37 |
| Colon cancer | 4 |
| Colorectal cancer | 252 |
| Corneal dystrophy | 1 |
| Coronary artery disease | 647 |
| Coronary atherosclerosis | 3 |
| Coronary heart disease | 1,471 |
| C-reactive protein measurement | 638 |
| Cutaneous melanoma | 72 |
| Deep vein thrombosis | 6 |
| Dementia | 23 |
| Diabetes mellitus | 6 |
| Diabetic retinopathy | 4 |
| Diverticulitis | 4 |
| Dupuytren contracture | 17 |
| Dysphagia | 1 |
| Emphysema | 22 |
| Endometriosis | 60 |
| End stage renal disease | 5 |
| Erectile dysfunction | 2 |
| Esophageal adenocarcinoma | 18 |
| Esophageal cancer | 21 |
| Essential tremor | 5 |
| Estrogen-receptor negative breast cancer | 39 |
| Estrogen-receptor positive breast cancer | 3 |
| Plantar Fasciitis | 1 |
| Fasting blood glucose measurement | 50 |
| Femoral neck bone mineral density | 56 |
| FEV ratio | 534 |
| Gastric carcinoma | 18 |
| Gastrointestinal hemorrhage | 1 |
| Glaucoma | 448 |
| Gout | 59 |
| Graves disease | 7 |
| HDL cholesterol change measurement | 8 |
| Heart failure | 18 |
| Heel bone mineral density | 1,262 |
| Hematuria | 2 |
| hepatocellular carcinoma | 11 |
| Herpes zoster | 7 |
| High density lipoprotein cholesterol measurement | 1,054 |
| Hip bone mineral density | 3 |
| Hyperplasia of prostate | 20 |
| Hypertension | 225 |
| Hypotension | 1 |
| Idiopathic dilated cardiomyopathy | 2 |
| Inflammatory bowel disease | 258 |
| Intracerebral hemorrhage | 5 |
| Ischemic Heart Disease | 53 |
| Ischemic stroke | 67 |
| Keratinocyte carcinoma | 206 |
| Kidney stone | 32 |
| Large artery stroke | 2 |
| Large cell lymphoma | 5 |
| Late-onset Alzheimers disease | 13 |
| LDL cholesterol change measurement | 12 |
| Lipoma | 29 |
| Low density lipoprotein cholesterol measurement | 717 |
| Lung adenocarcinoma | 42 |
| Lung cancer | 119 |
| Macular degeneration | 553 |
| Magnesium measurement | 7 |
| Malignant neoplasm | 1 |
| Mean platelet volume | 620 |
| Melanoma | 138 |
| Metabolic syndrome | 140 |
| Metabolite measurement | 806 |
| Mild cognitive impairment | 35 |
| Monoclonal gammopathy | 34 |
| Morbid obesity | 1 |
| Multiple myeloma | 37 |
| Multiple sclerosis | 346 |
| Myeloid leukemia | 12 |
| Myocardial infarction | 122 |
| Myopia | 510 |
| Nasal polyps | 13 |
| Neutropenia | 2 |
| Non-lobar intracerebral hemorrhage | 3 |
| Non-melanoma skin cancer | 41 |
| Non-small cell lung carcinoma | 52 |
| Obesity | 427 |
| Obstructive sleep apnea | 4 |
| Open-angle glaucoma | 253 |
| Orthostatic hypotension | 1 |
| Osteoarthritis | 126 |
| Osteoporosis | 1,258 |
| Ovarian cancer | 38 |
| Overweight body mass index status | 12 |
| Overweight | 75 |
| Paget's disease of bone | 8 |
| Pancreatic cancer | 29 |
| Parkinson's disease | 213 |
| Peripheral arterial disease | 8 |
| Primary angle-closure glaucoma | 7 |
| Primary open angle glaucoma | 2 |
| Prostate cancer | 266 |
| Proteinuria | 2 |
| Psoriasis | 118 |
| Psoriasis vulgaris | 26 |
| Psoriatic arthritis | 36 |
| QT interval | 116 |
| Renal cell carcinoma | 15 |
| Respiratory failure | 1 |
| Rheumatoid arthritis | 171 |
| Sjogren syndrome | 13 |
| Skin cancer | 42 |
| Sleep apnea | 4 |
| Small cell lung carcinoma | 3 |
| Spine bone mineral density | 75 |
| Squamous cell carcinoma | 66 |
| Squamous cell lung carcinoma | 23 |
| Sudden cardiac arrest | 3 |
| Systemic lupus erythematosus | 153 |
| Telomere length | 46 |
| Thyroid cancer | 112 |
| Triglyceride measurement | 900 |
| T-tau measurement | 9 |
| Type 2 diabetes | 1,833 |
| Type II diabetes mellitus | 1,559 |
| Urate measurement | 339 |
| Uric acid measurement | 128 |
| Varicose veins | 31 |
| Vascular dementia | 3 |
| Venous thromboembolism | 108 |
| Volumetric bone mineral density | 6 |
| Wet Macular degeneration | 8 |

**Table S3. Number of eQTL-eGene pairs associated with longevity variants in 47 GTEx tissues.** Tissues include 47 GTEx tissues; No. of eQTL-eGene pairs represents the total number of significant variant-gene associations in each tissue of GTEx v8; No. of longevity eQTL-eGene pairs represents the number of eQTL-eGene pairs associated with longevity variants in each tissue; No. of unique genes regulated by longevity variants is the number of unique genes their gene expression regulation associated with longevity variants. No. of unique longevity variants means the number of unique longevity variants that associated with gene expression in each tissue.

| Tissues | No. of eQTL-eGene pairs | No. of longevity eQTL-eGene pairs | No. of unique genes regulated by longevity variants | No. of unique longevity variants |
| --- | --- | --- | --- | --- |
| Adipose Subcutaneous | 2,930,371 | 92 | 60 | 30 |
| Adipose Visceral Omentum | 2,031,092 | 54 | 40 | 21 |
| Adrenal Gland | 959,377 | 25 | 16 | 17 |
| Artery Aorta | 2,038,600 | 63 | 39 | 23 |
| Artery Coronary | 721,638 | 22 | 17 | 9 |
| Artery Tibial | 2,958,255 | 72 | 49 | 24 |
| Brain Amygdala | 347,360 | 5 | 4 | 4 |
| Brain Anterior cingulate cortex BA24 | 531,519 | 10 | 6 | 8 |
| Brain Caudate basal ganglia | 906,717 | 26 | 16 | 12 |
| Brain Cerebellar Hemisphere | 1,183,392 | 24 | 18 | 13 |
| Brain Cerebellum | 1,505,828 | 28 | 18 | 16 |
| Brain Cortex | 1,062,869 | 24 | 14 | 13 |
| Brain Frontal Cortex BA9 | 770,795 | 12 | 9 | 7 |
| Brain Hippocampus | 541,494 | 11 | 6 | 7 |
| Brain Hypothalamus | 568,170 | 21 | 13 | 11 |
| Brain Nucleus accumbens basal ganglia | 908,867 | 26 | 15 | 18 |
| Brain Putamen basal ganglia | 695,348 | 21 | 14 | 11 |
| Brain Spinal cord cervical c-1 | 422,664 | 12 | 7 | 9 |
| Brain Substantia nigra | 283,775 | 6 | 3 | 6 |
| Breast Mammary Tissue | 1,616,204 | 43 | 35 | 18 |
| Colon Sigmoid | 1,474,958 | 42 | 28 | 18 |
| Colon Transverse | 1,672,710 | 43 | 32 | 16 |
| Esophagus Gastroesophageal Junction | 1,556,680 | 34 | 28 | 9 |
| Esophagus Mucosa | 2,616,829 | 82 | 48 | 36 |
| Esophagus Muscularis | 2,529,274 | 69 | 46 | 24 |
| Heart Atrial Appendage | 1,620,488 | 56 | 34 | 24 |
| Heart Left Ventricle | 1,409,425 | 46 | 32 | 14 |
| Kidney Cortex | 111,170 | 2 | 2 | 1 |
| Liver | 629,560 | 27 | 24 | 12 |
| Lung | 2,392,205 | 62 | 42 | 20 |
| Minor Salivary Gland | 422,972 | 9 | 7 | 6 |
| Muscle Skeletal | 2,601,576 | 83 | 49 | 29 |
| Nerve Tibial | 3,460,274 | 83 | 51 | 27 |
| Ovary | 570,917 | 4 | 4 | 3 |
| Pancreas | 1,340,449 | 37 | 29 | 13 |
| Pituitary | 1,226,884 | 27 | 18 | 13 |
| Prostate | 823,691 | 19 | 17 | 8 |
| Skin Not Sun Exposed Suprapubic | 2,697,338 | 85 | 57 | 37 |
| Skin Sun Exposed Lower leg | 3,244,343 | 89 | 62 | 39 |
| Small Intestine Terminal Ileum | 644,573 | 14 | 12 | 9 |
| Spleen | 1,262,025 | 25 | 22 | 9 |
| Stomach | 1,156,159 | 28 | 21 | 10 |
| Testis | 2,887,028 | 52 | 38 | 24 |
| Thyroid | 3,712,241 | 93 | 60 | 32 |
| Uterus | 328,521 | 5 | 5 | 3 |
| Vagina | 338,600 | 5 | 4 | 4 |
| Whole Blood | 2,414,654 | 75 | 53 | 22 |

**Table S4.** **GO functional annotation of up/down-regulated genes associated with longevity variants.** The first column shows the original database/resource where the terms orient. The second column shows the enriched terms associated with genes up-/down-regulated with age. The third column shows the Benjamini-Hochberg modified statistic for a p-value. Only selected top enriched terms with a p-value less than 5% are shown. Gene lists were classified into “up-regulated genes” and “down-regulated genes”.

| **Longevity allele associated with lower gene expression** | | | |
| --- | --- | --- | --- |
| **Category** | **Term** | | **Adjusted p-value** |
| GOTERM_CC_FAT | GO:0042611~MHC protein complex | | 9.19E-09 |
| GOTERM_CC_FAT | GO:0071556~integral component of lumenal side of endoplasmic reticulum membrane | | 1.85E-07 |
| GOTERM_CC_FAT | GO:0098553~lumenal side of endoplasmic reticulum membrane | | 1.85E-07 |
| GOTERM_CC_FAT | GO:0012507~ER to Golgi transport vesicle membrane | | 2.40E-07 |
| INTERPRO | IPR003006:Immunoglobulin/major histocompatibility complex, conserved site | | 3.23E-07 |
| KEGG_PATHWAY | hsa05332:Graft-versus-host disease | | 9.21E-07 |
| KEGG_PATHWAY | hsa05330:Allograft rejection | | 9.53E-07 |
| KEGG_PATHWAY | hsa04940:Type I diabetes mellitus | | 1.41E-06 |
| GOTERM_CC_FAT | GO:0042613~MHC class II protein complex | | 1.60E-06 |
| GOTERM_CC_FAT | GO:0030134~ER to Golgi transport vesicle | | 1.69E-06 |
| KEGG_PATHWAY | hsa05320:Autoimmune thyroid disease | | 3.95E-06 |
| KEGG_PATHWAY | hsa05416:Viral myocarditis | | 5.53E-06 |
| GOTERM_CC_FAT | GO:0030658~transport vesicle membrane | | 1.29E-05 |
| KEGG_PATHWAY | hsa04612:Antigen processing and presentation | | 2.59E-05 |
| GOTERM_CC_FAT | GO:0030133~transport vesicle | | 6.18E-05 |
| GOTERM_CC_FAT | GO:0030662~coated vesicle membrane | | 6.25E-05 |
| GOTERM_CC_FAT | GO:0098552~side of membrane | | 6.41E-05 |
| GOTERM_CC_FAT | GO:0030176~integral component of endoplasmic reticulum membrane | | 7.08E-05 |
| GOTERM_MF_FAT | GO:0032395~MHC class II receptor activity | | 1.15E-04 |
| KEGG_PATHWAY | hsa05168:Herpes simplex infection | | 3.57E-04 |
| GOTERM_BP_FAT | GO:0002478~antigen processing and presentation of exogenous peptide antigen | | 3.47E-04 |
| GOTERM_BP_FAT | GO:0019884~antigen processing and presentation of exogenous antigen | | 3.47E-04 |
| GOTERM_BP_FAT | GO:0048002~antigen processing and presentation of peptide antigen | | 3.58E-04 |
| GOTERM_BP_FAT | GO:0060333~interferon-gamma-mediated signaling pathway | | 3.83E-04 |
| KEGG_PATHWAY | hsa04514:Cell adhesion molecules (CAMs) | | 7.16E-04 |
| KEGG_PATHWAY | hsa04145:Phagosome | | 8.63E-04 |
| GOTERM_CC_FAT | GO:0030666~endocytic vesicle membrane | | 9.63E-04 |
| GOTERM_MF_FAT | GO:0042605~peptide antigen binding | | 8.32E-04 |
| KEGG_PATHWAY | hsa05150:Staphylococcus aureus infection | | 0.0012308 |
| GOTERM_BP_FAT | GO:0019882~antigen processing and presentation | | 0.00128071 |
| GOTERM_CC_FAT | GO:0030139~endocytic vesicle | | 0.00201026 |
| KEGG_PATHWAY | hsa05169:Epstein-Barr virus infection | | 0.00258542 |
| GOTERM_CC_FAT | GO:0000139~Golgi membrane | | 0.00266798 |
| KEGG_PATHWAY | hsa05310:Asthma | | 0.00308433 |
| UP_SEQ_FEATURE | region of interest:Alpha-1 | | 0.00251383 |
| UP_SEQ_FEATURE | region of interest:Alpha-2 | | 0.00251383 |
| GOTERM_CC_FAT | GO:0030659~cytoplasmic vesicle membrane | | 0.00347482 |
| GOTERM_CC_FAT | GO:0044433~cytoplasmic vesicle part | | 0.00376096 |
| GOTERM_CC_FAT | GO:0012506~vesicle membrane | | 0.00394322 |
| GOTERM_BP_FAT | GO:0071346~cellular response to interferon-gamma | | 0.00538635 |
| **Longevity allele associated with higher gene expression** | | | |
| **Category** | **Term** | **Adjusted p-value** | |
| OMIM_DISEASE | Six new loci associated with blood low-density lipoprotein cholesterol, high-density lipoprotein cholesterol or triglycerides in humans | 0.0010346 | |
| OMIM_DISEASE | Newly identified loci that influence lipid concentrations and risk of coronary artery disease | 0.0010796 | |
| OMIM_DISEASE | Common variants at 30 loci contribute to polygenic dyslipidemia | 0.00212591 | |
| OMIM_DISEASE | LDL-cholesterol concentrations: a genome-wide association study | 0.00292168 | |
| GOTERM_BP_FAT | GO:0010033~response to organic substance | 0.01793415 | |
| GOTERM_BP_FAT | GO:0043279~response to alkaloid | 0.02155777 | |
| GOTERM_BP_FAT | GO:0035094~response to nicotine | 0.02881208 | |

**Table S5.** **Statistics for comparison of eQTL-eGene pairs between GTEx and previous independent studies.**

Here we performed a comparison of the eQTL-eGene pairs from GTEx study to prior six independent studies in five tissues (subcutaneous fat, brain cortex, heart left ventricle, lung and whole blood). No. of GTEx pairs defers to the number of eQTL-eGene pairs generated from GTEx in each tissue; No. of reported pairs is the number of eQTL-eGene pairs from independent studies. No. of overlapped eQTL is the number of overlapped eQTL between previous studies and GTEx; No. of overlapped pairs is the number of overlapped eQTL-eGene pairs between independent studies and GTEx; No. of same expr dir is the number of target allele regulate the gene expression in the same direction between GTEx and other studies; No. of diff expr dir is the number of target allele regulate the gene expression different between GTEx and other studies.

| Tissue |  | MuTHER Fat | Rosmap Brain | Heart | Lung | Blood1 | Blood2 |
| --- | --- | --- | --- | --- | --- | --- | --- |
| Subcutaneous Fat | No. of GTEx pairs | 2,930,371 | | | | | |
|  | No. of reported pairs | 22,989 | 373,463 | 6,081 | 15,231 | 853,485 | 10,506,072 |
|  | No. of overlapped eQTL | 8,562 | 292,235 | 4,985 | 8,604 | 467,624 | 4,744,631 |
|  | No. of overlapped pairs | 2,347 | 171,473 | 2,782 | 3,946 | 128,651 | 1,196,976 |
|  | No. of same expr dir | 2,192 | 154,544 | 2,622 | 3,578 | 104,965 | 1,073,253 |
|  | No. of diff expr dir | 155 | 16,929 | 160 | 358 | 23,686 | 123,723 |
| Brain Cortex | No. of GTEx pairs | 1,062,869 | | | | | |
|  | No. of reported pairs | 22,989 | 373,463 | 6,081 | 15,231 | 853,485 | 10,506,072 |
|  | No. of overlapped eQTL | 4,154 | 261,872 | 4,179 | 4,716 | 255,945 | 2,555,613 |
|  | No. of overlapped pairs | 893 | 167,005 | 2,258 | 1,725 | 49,042 | 409,053 |
|  | No. of same expr dir | 770 | 164,836 | 2,042 | 1,499 | 37,475 | 346,119 |
|  | No. of diff expr dir | 123 | 2,169 | 216 | 226 | 11,567 | 62,934 |
| Heart Left Ventricle | No. of GTEx pairs | 1,409,425 | | | | | |
|  | No. of reported pairs | 22,989 | 373,463 | 6,081 | 15,231 | 853,485 | 10,506,072 |
|  | No. of overlapped eQTL | 5,422 | 248,730 | 4,968 | 5,995 | 317,465 | 3,153,043 |
|  | No. of overlapped pairs | 1,308 | 136,187 | 3,143 | 2,496 | 71,469 | 609,471 |
|  | No. of same expr dir | 1,180 | 128,592 | 3,003 | 2,246 | 57,330 | 528,843 |
|  | No. of diff expr dir | 128 | 7,595 | 140 | 250 | 14,139 | 80,628 |
| Lung | No. of GTEx pairs | 2,392,205 | | | | | |
|  | No. of reported pairs | 22,989 | 373,463 | 6,081 | 15,231 | 853,485 | 10,506,072 |
|  | No. of overlapped eQTL | 7,234 | 279,215 | 4,803 | 8,548 | 422,226 | 4,246,023 |
|  | No. of overlapped pairs | 1,681 | 156,741 | 2,774 | 4,278 | 111,436 | 990,738 |
|  | No. of same expr dir | 1,500 | 141,594 | 2,551 | 4,034 | 94,747 | 916,119 |
|  | No. of diff expr dir | 181 | 15,147 | 223 | 244 | 16,689 | 74,619 |
| Whole Blood | No. of GTEx pairs | 2,414,654 | | | | | |
|  | No. of reported pairs | 22,989 | 373,463 | 6,081 | 15,231 | 853,485 | 10,506,072 |
|  | No. of overlapped eQTL | 7,382 | 268,251 | 4,749 | 7,509 | 477,240 | 4,651,543 |
|  | No. of overlapped pairs | 1,583 | 136,368 | 2,617 | 3,170 | 169,509 | 1,362,748 |
|  | No. of same expr dir | 1,303 | 116,102 | 2,301 | 2,757 | 158, 591 | 1,333,290 |
|  | No. of diff expr dir | 280 | 20,266 | 316 | 413 | 10,918 | 29,458 |

**Table S6. Statistics for comparison of disease/longevity-associated eQTL-eGene pairs from GTEx and previous independent studies.**

Here we performed a comparison of the disease/longevity-associated eQTL-eGene pairs from GTEx study to prior independent studies in five tissues (subcutaneous fat, brain cortex, heart left ventricle, lung and whole blood). No. of disease/longevity eQTL defers to the number of GTEx eGenes are regulated by longevity and disease SNPs in each tissue. No. of reported eQTL is the number of eQTL-eGene pairs from previous studies. No. of overlapped eQTL is the number of overlapped eQTL-eGene pairs between previous studies and GTEx, number in the bracket is the unique number of variant-gene-variant trio.

| Tissue |  | MuTHER Fat | Rosmap Brain | Heart | Lung | Blood1 | Blood2 |
| --- | --- | --- | --- | --- | --- | --- | --- |
| Subcutaneous Fat | No. of disease/longevity eQTL | 322 | | | | | |
|  | No. of duplicated pairs | 2,166 | 154,506 | 2,026 | 3,093 | 104,876 | 1,072,131 |
|  | No. of overlapped eQTL | 0 | 41(18) | 4(3) | 4(3) | 97(71) | 574(427) |
| Brain Cortex | No. of disease/longevity eQTL | 55 | | | | | |
|  | No. of duplicated pairs | 761 | 164,724 | 1,591 | 1,273 | 37,467 | 345,640 |
|  | No. of overlapped eQTL | 0 | 30(8) | 0 | 0 | 54(40) | 169(116) |
| Heart Left Ventricle | No. of disease/longevity eQTL | 129 | | | | | |
|  | No. of duplicated pairs | 1,169 | 128,567 | 2,340 | 1,928 | 57,299 | 528,587 |
|  | No. of overlapped eQTL | 0 | 33(11) | 2(1) | 1(1) | 69(50) | 316(241) |
| Lung | No. of disease/longevity eQTL | 190 | | | | | |
|  | No. of duplicated pairs | 1,485 | 141,457 | 1,958 | 3,514 | 94,708 | 915,467 |
|  | No. of overlapped eQTL | 0 | 4(4) | 2(1) | 4(3) | 19(14) | 275(227) |
| Whole Blood | No. of disease/longevity eQTL | 257 | | | | | |
|  | No. of duplicated pairs | 1,286 | 116,090 | 1,733 | 2,358 | 158,445 | 1,331,137 |
|  | No. of overlapped eQTL | 1(1) | 34(12) | 2(1) | 2(1) | 142(107) | 609(479) |

**Table S7. Number of disease/longevity alleles association with gene expression regulation.**

The first column shows the number of gene expression direction associated with disease alleles are different from longevity alleles; The second column shows the number of unique variant-gene-variant trios; eGenes is the number of unique genes which their gene expression associated with disease/longevity variants

| Tissue | Gene exp dir of disease allele != longevity allele (eGene) | Gene exp dir of disease allele == longevity allele (eGene) |
| --- | --- | --- |
| Subcutaneous Fat | 323 (33) | 132 (30) |
| Brain Cortex | 103 (6) | 17 (4) |
| Heart Left Ventricle | 187 (15) | 63 (14) |
| Lung | 150 (23) | 85 (19) |
| Whole Blood | 397 (35) | 106 (30) |
